# Comparative analysis of the molecular mechanism of resistance to vapendavir across a panel of picornavirus species

**DOI:** 10.1101/2020.07.16.207373

**Authors:** Kristina Lanko, Liang Sun, Mathy Froeyen, Pieter Leyssen, Leen Delang, Carmen Mirabelli, Johan Neyts

## Abstract

Vapendavir is a rhino/enterovirus inhibitor that targets a hydrophobic pocket in the viral capsid. Drug-resistant variants were selected *in vitro*. Mutations in the drug-binding pocket in VP1 (C199R/Y in hRV14; I194F in PV1; M252L and A156T in EV-D68), typical for this class of compounds, were identified. We also observed mutations that are located outside the pocket (K167E in EV-D68 and G149C in hRV2) and that contribute to the resistant phenotype. Remarkably, the G149C substitution made the replication of human rhinovirus 2 dependent on the presence of vapendavir. Our data suggest that vapendavir binding to the capsid of the dependent isolate may be required to stabilize the viral particle and to allow efficient dissemination of the virus. Our results demonstrate that vapendavir-resistant pheno- and genotypes of clinically relevant picornavirus species are more complex than generally believed.

## Introduction

The *Picornaviridae* family comprises numerous human pathogens, belonging to species Enterovirus A-D and Rhinoviruses A-C. Clinical manifestations of picornavirus infections range from mild (e.g. common cold caused by Rhinoviruses (hRV); hand-foot-and-mouth disease by coxsackievirus A (CVA), enterovirus 71 (EV-A71) and others) to life-threatening (myocarditis, encephalitis, acute flaccid paralysis), although many infections remain subclinical with no apparent symptoms [1].

Human rhinoviruses are the major causative agents of common cold affecting millions of people worldwide [2]. Rhinoviruses have been shown to not only cause mild upper respiratory disease but also to provoke the exacerbations of preexisting respiratory conditions like COPD and asthma [3,4]. EV-D68 recently emerged as another picornavirus with respiratory tropism [5]. In addition, during recent outbreaks of EV-D68 the link between this viral infection and neurological complications (acute flaccid myelitis, AFM) has become clear [6–10]. Despite the great success of the Global Polio Eradication Initiative (GPEI), poliovirus (PV) will remain a threat even after complete eradication of the wild-type viruses, e.g. due to circulation of vaccine-derived viruses or accidental virus release[11]. Development of antiviral drugs to control and prevent such potential outbreaks is one of the priorities in the polio endgame [12].

To date there is no specific antiviral therapy for any of the picornavirus infections. Vaccines are only available for poliovirus (live-attenuated oral polio vaccine and inactivated vaccine) [13] and EV-A71 (approved in China) [14]. Several classes of early stage inhibitors – capsid binders – have been developed, of which some entered clinical trials [12,15,16]. WIN-like compounds such as pirodavir, pleconaril, pocapavir and vapendavir bind to the pocket located in the structural protein VP1 in the canyon – a structure present in most picornaviruses [17]. This region plays a major role in early stages of picornavirus infections (attachment, entry and uncoating). Capsid binders increase the stability of the viral capsid, thus preventing conformational changes necessary for receptor interaction and uncoating. Vapendavir has initially been developed as an hRV inhibitor and has progressed to phase IIb clinical trials; the drug results in a reduction of viral load in hRV infected adults. However, it failed to reduce asthma exacerbations [18] (ClinicalTrials.gov: NCT02367313). Like other capsid binders, vapendavir exerts broad-spectrum activity against different enteroviruses including EV-A71, EV-D68, PV, and human rhinoviruses [19–22]. One of the drawbacks of capsid binders is the rapid development of resistance *in vitro* [20,21,23,24] and in men [12,15]. Typical mutations that have been reported to be associated with resistance to capsid binders are I99F in hRV2, A150T/V, C199R, V188I and E276K in hRV14; V69A and K155E in EV-D68; A24V and I194F/M in PV1. The selection of vapendavir-resistant isolates has not been studied yet, however the cross-resistance of pleconaril-resistance rhinoviruses to vapendavir [21] suggests similar mechanisms of virus adaptation. Here, we explore the particular characteristics of resistance development of four enterovirus species (hRV14, hRV2, EV-D68 and PV) to vapendavir.

## Materials and methods

### Cells and viruses

All cell types used in this study [Buffalo green monkey (BGM) cells (ECACC 90092601), Human Caucasian embryo rhabdomyosarcoma (RD, ECACC 85111502), Epithelium cells cervix adenoncarcinoma (HeLa) kindly provided by Dr. K. Andries (Janssen Pharmaceutica, Belgium)] were cultured in MEM Rega3 medium (Gibco) supplemented with 10% FBS, 2mM L-glutamine (Gibco) and 0.075% sodium bicarbonate (Gibco) at 37°C in 5% CO_2_ incubators.

Viral infections were carried out in the same medium supplemented with 2% FBS. hRV and EV-D68 infections were carried out in the culture medium which was supplemented with 30 mM Mg^2+^ at 35°C in 5% CO_2_ incubators.

PV1 (Sabin) was derived from the infectious clone pT7/S1F, which was kindly provided by A.J. Macadam [25] and virus stocks were produced in BGM cells. The EV-D68 strain CU70 was obtained from RIVM (Bilthoven, The Netherlands) and cultured in HeLa cells. hRV2 and hRV14 were kindly provided by Dr. K. Andries (Janssen Pharmaceutica, Belgium) and cultured in HeLa cells.

### Compounds

Vapendavir (BTA798) was provided by Vaxart Inc., USA. Pirodavir was synthesized by Prof. G. Pürstinger (University of Innsbruck). Pleconaril was kindly provided by V. Makarov (RAS Institute of Biochemistry, Russia). Compound stocks were prepared in DMSO at a concentration of 10 mM and diluted in cell culture medium for experimental use.

### 5 step resistance selection protocol

The detailed description of the protocol has been published previously [26]. In brief, optimal concentrations of vapendavir and virus input were determined for each virus. Optimal conditions are defined as the lowest compound concentration and highest virus input still resulting in complete inhibition of viral-induced CPE. Three identical 96-well plates were set up with the optimal condition and development of CPE was monitored microscopically. The supernatant from the wells exhibiting CPE (potentially resistant virus) was passaged in the presence of vapendavir to obtain a virus stock and the sensitivity of this stock was tested in an antiviral assay.

### Antiviral assays

Cells were seeded in 96-well plates (HeLa 15*10^3^ cells/well; BGM 20*10^3^ cells/well) in 2% FBS containing MEM Rega3 medium and were kept overnight at 37°C or 35°C (for hRV) 5% CO_2_ to allow attachment. The next day serial dilutions of the compounds were added to the cells. Cultures were infected with viruses at a MOI 0,01 and were kept until full CPE was observed in virus control (24h for PV1-Sabin, 3 days for other viruses in this study). Microscopic evaluation and cell viability readout with MTS reagent [3-(4,5-dimethylthiazol-2-yl)-5-(3-carboxymethoxyphenol)-2-(4-sulfophenyl)-2H-tetrazolium, inner salt; Promega] were performed for assessing the activity of compounds.

### Replication kinetics

HeLa cells were seeded in 48-well plates and infected with MOI 2 for 1h at 35°C. After removal of the inoculum, cultures were washed 3 times with PBS and fresh medium was added. Intracellular RNA was isolated at indicated timepoints with the RNeasy mini kit (Qiagen) and quantified by qRT-PCR.

### Thermostability assay

HRV2 WT or C3 isolate stocks were incubated at 37°C for 15 min and then 2 min at temperatures between 37-62°C (5°C increment) in presence of 5 μM vapendavir or DMSO solvent. The samples were cooled down to 4°C and then the infectivity was determined by end-point titration.

### qRT-PCR

qRT-PCR was performed using the BioRad iTaq Universal SYBR Green kit with pan-enterovirus primers ENRI 4- and ENRI 3+ [27] on Applied Biosystems 7500 Fast Real-Time PCR System. A geneBlock (IDT) standard corresponding to the target region of the primers (5’UTR) was included for quantification of viral RNA copies.

### Sequencing

Viral RNA was isolated from virus stocks with Nucleospin RNA virus (Macherey-Nagel) according to manufacturer protocol. CDNA fragments were amplified with strain-specific primers using the one step RT-PCR kit (Qiagen). Sequencing was performed by Macrogen Europe. VectorNTI and SnapGene software were used for analysis.

### Site-directed mutagenesis

Mutagenesis was performed on the plasmids pT7/S1F PV1 [28], pT7/EV-A71BrCr and pCMV/hRV14 by using the QuikChange II XL Site-Directed Mutagenesis Kit (Agilent Technologies). All mutants were verified by sequencing after mutagenesis. RNA was obtained by using a T7 RiboMAX Large Scale RNA Production System (Promega) and infectious viruses were generated by transfecting RNA into respective cells with the TransIT-mRNA Transfection Kit (Mirus). pCMV/hRV14 plasmid was transfected with TransIT-LT1 Transfection Reagent (Mirus) into HeLa cells.

### Molecular modelling

A Molecular dynamics simulation with explicit water molecules was performed using the Amber18 software[29] for the WT pentamer (PDB entry 3vdd [21])and a pentamer where all five g149.a were mutated to cysteine. 4 simulations were setup starting from the pentamer structure in pdb entry 3vdd: a wt pentamer with and without 5 vapendavir molecules bound; the g149c.a pentamer with and without 5 vapendavir inhibitors. A total of 60 ns was simulated for all systems. We obtained stable simulations for all systems. Details of the simulation parameters are given in table S2. The LIGPLOT program was used to inspect the interactions in pentamer-vapendavir complexes[30]. Visualization of structures was performed with UCSF Chimera [31].

## Results

### Vapendavir-resistant variants are readily selected in cell culture

To obtain vapendavir-resistant variants of hRV14, hRV2, EV-D68 (CU70) and PV1 (Sabin) a 5-step resistance selection protocol was employed [26]. For each virus, the optimal resistance selection conditions were determined and three 96-well plates were set up for selection. We were able to obtain up to 7 potentially resistant populations for each virus. Virus stocks of these isolates were generated in further selection steps and the activity of vapendavir against these isolates was assessed in a multicycle CPE-reduction assay. Seven (7) resistant isolates of hRV14, 3 of PV1_Sabin, 1 of hRV2, and 2 of EV-D68 (CU70) were obtained. All the isolates were fully resistant against vapendavir (Table 1).

**Table 1.** Vapendavir-resistant variants carry mutations in VP1 and are cross-resistant to other capsid binders. Antiviral activity of capsid binders was assessed in a CPE-reduction assay. Data are mean values of 3 independent experiments ±SD. NA – not active

| Virus | Genotype VP1 | EC<sub>50</sub> μM (fold change EC<sub>50</sub>) | | |
| --- | --- | --- | --- | --- |
|  |  | Vapendavir | Pleconaril | Pirodavir |
| hRV14 | WT | 0.09 ± 0.01 | 0.20 ± 0.005 | 0.14 ± 0.1 |
| | C199R | >10 (>110) | 2.9 ± 1.0 (14.5) | >27 (>192) |
| | C199Y | >10 (>110) | 3.4 ± 1.6 (17) | 8.5 ± 0.5 (60) |
| hRV2 | WT | 0.04 ± 0.003 | 0.44 ± 0.1 | 0.46 ± 0.2 |
|  | G149C | >10 (>250) | >26 (>59) | >27 (>58) |
| PV1_Sabin | WT | 2.6 ± 0.04 | NA | 33 ± 1 |
|  | I194F | >10 (>3.8) | NA | >54 (>1.6) |
| EVD68_CU70 | WT | 1.4 ± 0.5 | 0.13 ± 0.06 | 2.4 ± 0.3 |
| | K167E_M252L | > 10 (>7) | 2.9 ± 1.6 (22) | > 54 (>22) |
| | A156T_K167E | > 10 (>7) | 1.4 ± 0.3 (10) | > 54 (>22) |

### Mutations in VP1 confer the capsid binder-resistant phenotype

We next sequenced the capsid region of the resistant isolates and identified amino acid substitutions in the VP1 protein as expected (Table 1). The resistant variants of hRV2, hRV14 and PV1 (Sabin) carried single amino acid substitutions in the VP1 region, whereas EV-D68 (CU70) variants carried double mutations with K167E present in both isolates. Reverse-engineering of the mutations identified in hRV14 and PV1 (Sabin) was performed and revealed that the introduction of respective mutations into the WT backbone resulted in a resistant phenotype (Table S1). To study whether the vapendavir-resistant isolates are cross-resistant against other capsid binders, the activity of two other capsid binders, pleconaril and pirodavir, against the vapendavir-resistant isolates was assessed. Cross-resistance to both compounds was observed for all isolates (Table 1).

### Location of mutations in VP1 structure

To determine the location of the identified substitutions in VP1, we superimposed available VP1 crystal structures of the virus strains used in our study and marked the location of mutated residues on the structure of hRV2 in complex with vapendavir PDB: 3vdd (Fig. 1, Fig. S1)[21]. The substitutions in the resistant populations of hRV14 and PV1 (Sabin) are located in the well-known drug-binding pocket [28,32], whereas the substitution in hRV2 is located outside the pocket in one of the VP1 loops not directly interacting with the compound. The K167E residue identified in both isolates of EV-D68 is located on the same loop outside the drug-binding pocket as in hRV2; however, two other residues (A156T and M252L) observed in EV-D68 resistant isolates are present in the drug-binding pocket.

**Figure 1.**
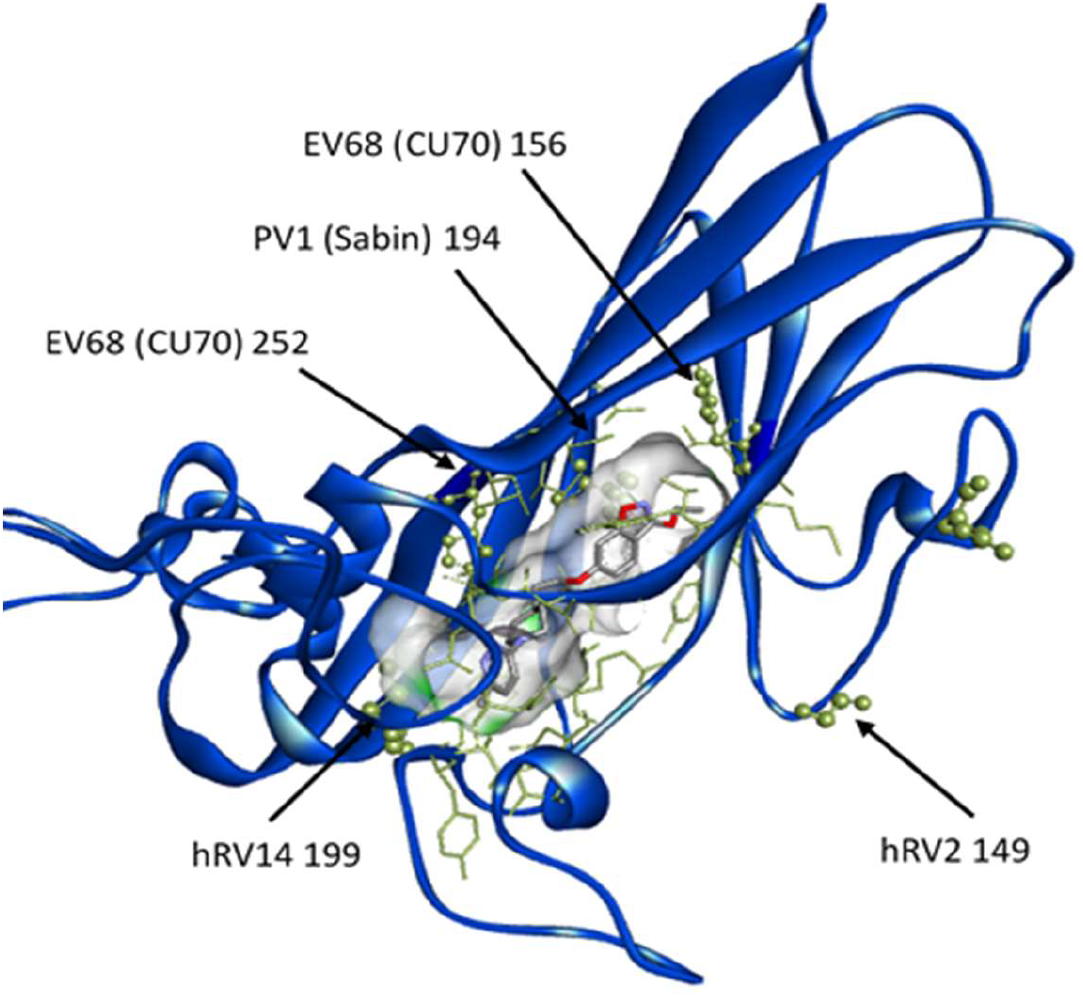
Location of identified mutated residues corresponding to the residues in hRV2 VP1 protein in complex with Vapendavir. VP1 protein in blue. Pocket surface in gray with drug-interacting residues in yellow. Residues mutated in resistant populations represented in sticks and balls.

### The infectivity of the hRV2 vapendavir-resistant isolate is dependent on vapendavir

The hRV2 vapendavir-resistant isolate C3 (hRV2_C3 carrying G149C) has a particular phenotype that was not observed in other resistant isolates that we obtained. The infectious virus titers were much lower for this mutant compared to WT hRV2, which was not the case for the resistant isolates of hRV14 and EV-D68 (Fig.S2). Much higher infectious titers were obtained when hRV2_C3 was titrated in the presence of 1 μg/mL of either vapendavir, pleconaril or pirodavir (Fig. 2A). The hRV2_C3 populations from the cultures that exhibited CPE at the highest dilution of the virus stock (1/100 without compound and 1/1000 in presence of vapendavir) were sequenced. Virus cultured in the absence of the drug lost the G149C mutation.

**Figure 2.**
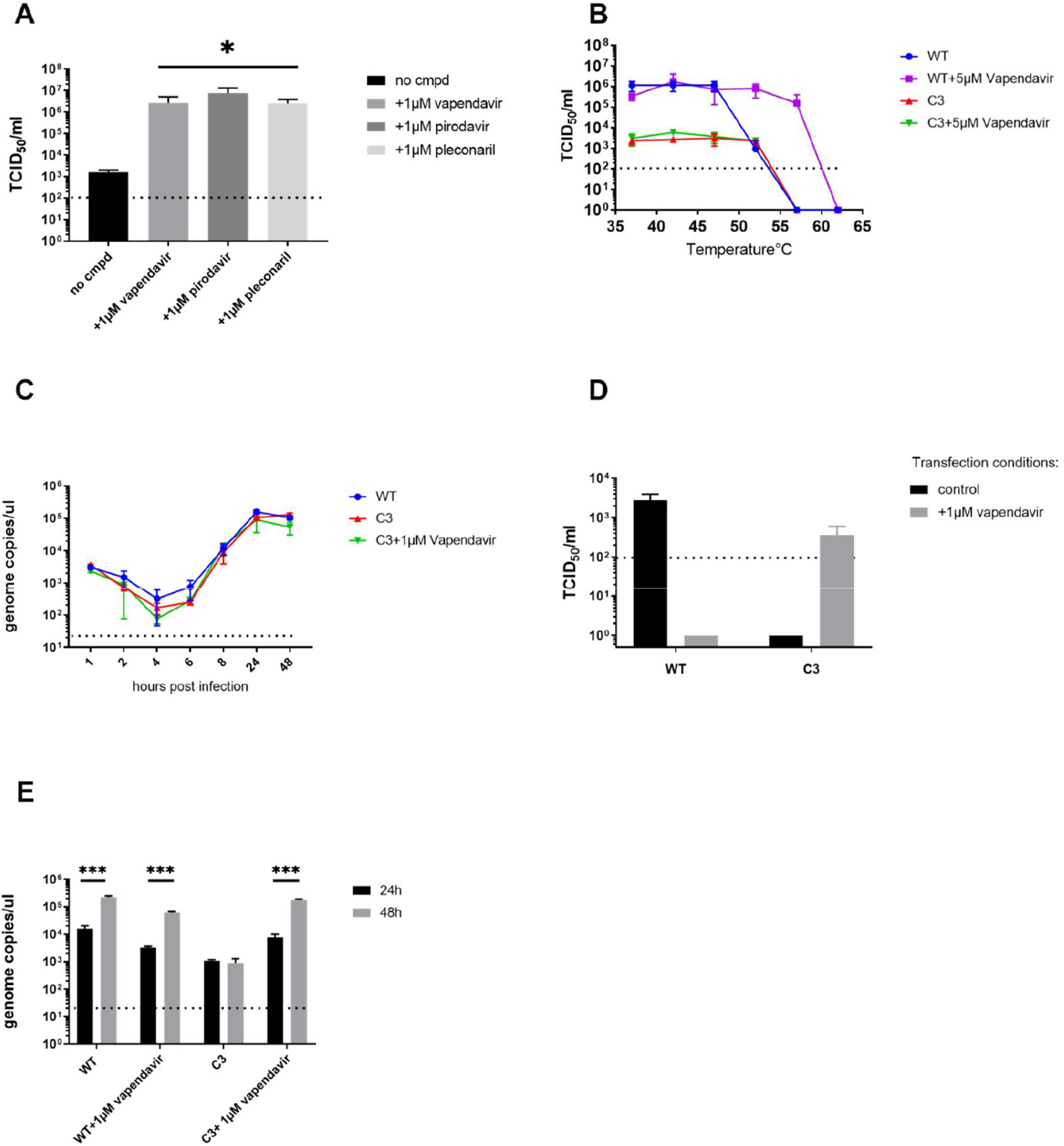
The replication of hRV2_C3 variant is dependent on Vapendavir. (A) Quantification of infectious hRV2_C3 virus grown in presence of different capsid binders. Data are mean values of 3 independent experiments ±SD. *, p< 0.05 (t-test) (B) Thermostability test of hRV2 WT and resistant variant C3. Data are mean values of 3 independent experiments ±SD (C) Replication kinetics of WT and mutant hRV2 follow the same pattern as assessed by qRT-PCR of intracellular viral RNA levels. Data are mean values of 2 independent experiments ±SD (D) Detection of infectious virus in culture supernatants at 24h post transfection by end-point titration in presence (isolate C3) or absence of vapendavir (WT virus). Transfection of viral RNA was performed either without (black bars) or with vapendavir (grey bars). Data are mean values of 3 independent experiments ±SD (E) Quantification of viral RNA in culture supernatants at 24h and 48h post transfection. ***, p<0.0005, analyzed by t-test. Data are mean values of 3 independent experiments ±SD.

Next viral RNA of the C3 mutant and WT were transfected in cells to test whether infectious virions can be produced in presence or absence of vapendavir (Fig. 2D). No infectious C3 virus was not detectable in the culture supernatant at 24h and 48h post transfection in the absence of antiviral pressure. However, when transfection of cultures with C3 RNA was performed in medium that was supplemented with 1 μM of vapendavir, infectious virus was obtained as could be detected by end-point titration in vapendavir-containing medium. This was not the case in the absence of the compound. Thus, the infectivity of C3 depends on vapendavir. There was also no increase in viral RNA copies from 24h to 48h in C3 transfected cultures in which no vapendavir was present in the medium. By contrast, when 1 μM of vapendavir was present in the transfected culture, there was a 1 log_10_ increase in viral genome copies from 24h to 48h as compared to untreated control, confirming the dissemination of the C3 virus in presence of the compound (Fig. 2E).

### Vapendavir does not protect the hRV2_C3 resistant mutant against heat-inactivation

The binding of WIN-like compounds to the viral capsid is known to make the virus less sensitive to heat-inactivation [33,34]. It has been previously reported that the mutations in the drug-binding pocket abolish the interaction between the capsid and the compound and as a consequence no protection against heat-inactivation is observed. To explore whether this type of interaction is possible with hRV2_C3, we performed a thermostability test (Fig.2B). The WT virus was inactivated at 56°C in the absence of compound and in presence of 5 μM of vapendavir at 62°C. This shift was not observed with the resistant isolate, indicating that vapendavir has no influence on heat-inactivation of the virus. In addition, there was no decrease in infectivity of the resistant isolate at 52°C as was observed for the WT virus. HRV2_C3 has however, the same plaque as the WT (data not shown), also the replication kinetics of the isolate and WT virus are comparable (Fig. 2C).

### The conserved glycine at VP1 position 149 is important for enterovirus infectivity

Since the G149 residue is conserved in many enterovirus species (Fig.S3), we introduced the G to C (G156C in RV14, G159C in EV-A71) substitution in infectious clones of hRV14 and EV-A71 BrCr and determined whether this would result in a similar resistant/dependent phenotype. Following transfection of the mutated hRV14 or EV-A71 infectious clone in HeLa cells, no infectious virus was detectable either in the presence or in absence of vapendavir. However, when the supernatant from the cultures transfected with EV-A71 G159C was passaged in absence of vapendavir, the virus reverted to WT and CPE was observed. No resistance or dependency on vapendavir was thus observed with the G to C substitution in EV-A71 or hRV14 background.

### Molecular modelling of the resistance mechanism of hRV2 isolate to vapendavir

Molecular modelling was used to explore whether the G149C in hRV2 affects the inhibitor binding in the pocket via network interactions (Fig. 3). The ligplot analysis [30] of 3vdd.pdb depicting the inhibitor interactions with surrounding amino acids (Fig. 4) reveals that M213 is one of the amino acids that may stabilize the binding of vapendavir by van der Waals (vdw) interactions. A contact analysis from the MD trajectory of the unliganded mutant simulation shows that there are many (vdw) contacts between the sidechains of C149 and N212 (Table S3). This is also clear from a representative structure from the MD simulation of the mutant with vapendavir (Fig. 5). Next, the Root mean square deviations (RMSD) and fluctuations (RMSF) for the 2 loops of amino acids 210-214 (containing M213, green ribbon in figure 5) and amino acids 147-151 (containing the G to C149 mutation, cyan ribbon in figure 5) in both WT and mutant (liganded and unliganded) simulations were explored. However, no significant differences between wildtype and mutant were observed.

**Figure 3.**
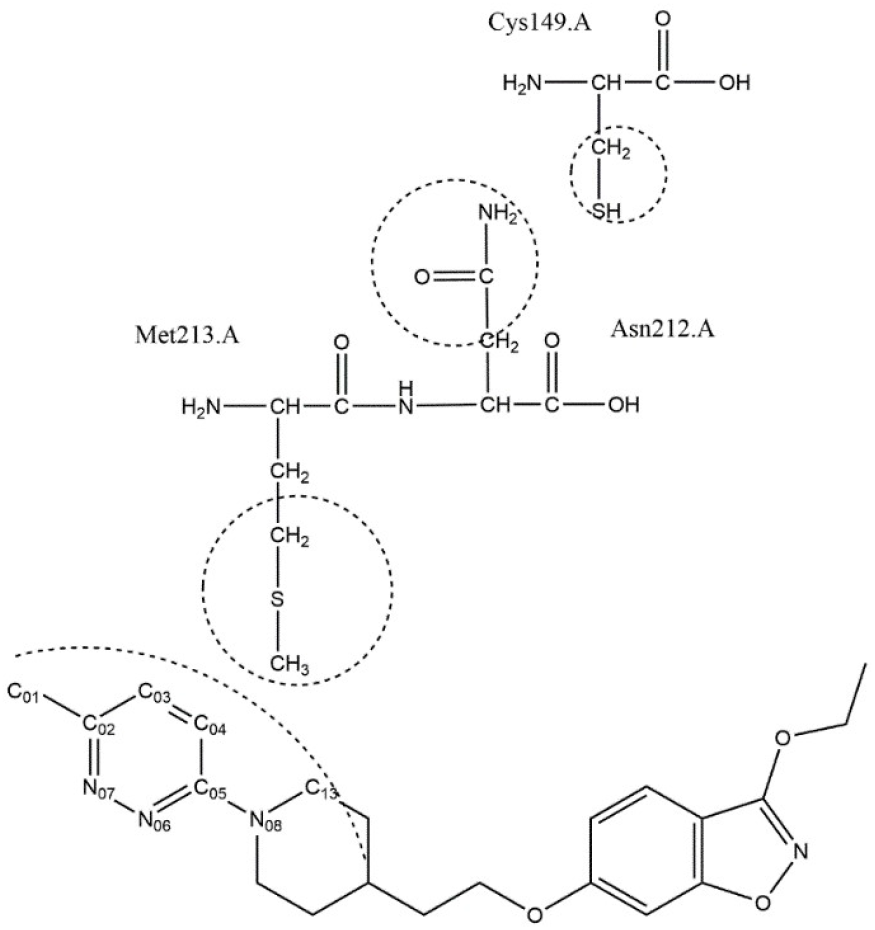
Vapendavir interaction map with VP1 residues of hRV2.

**Figure 4.**
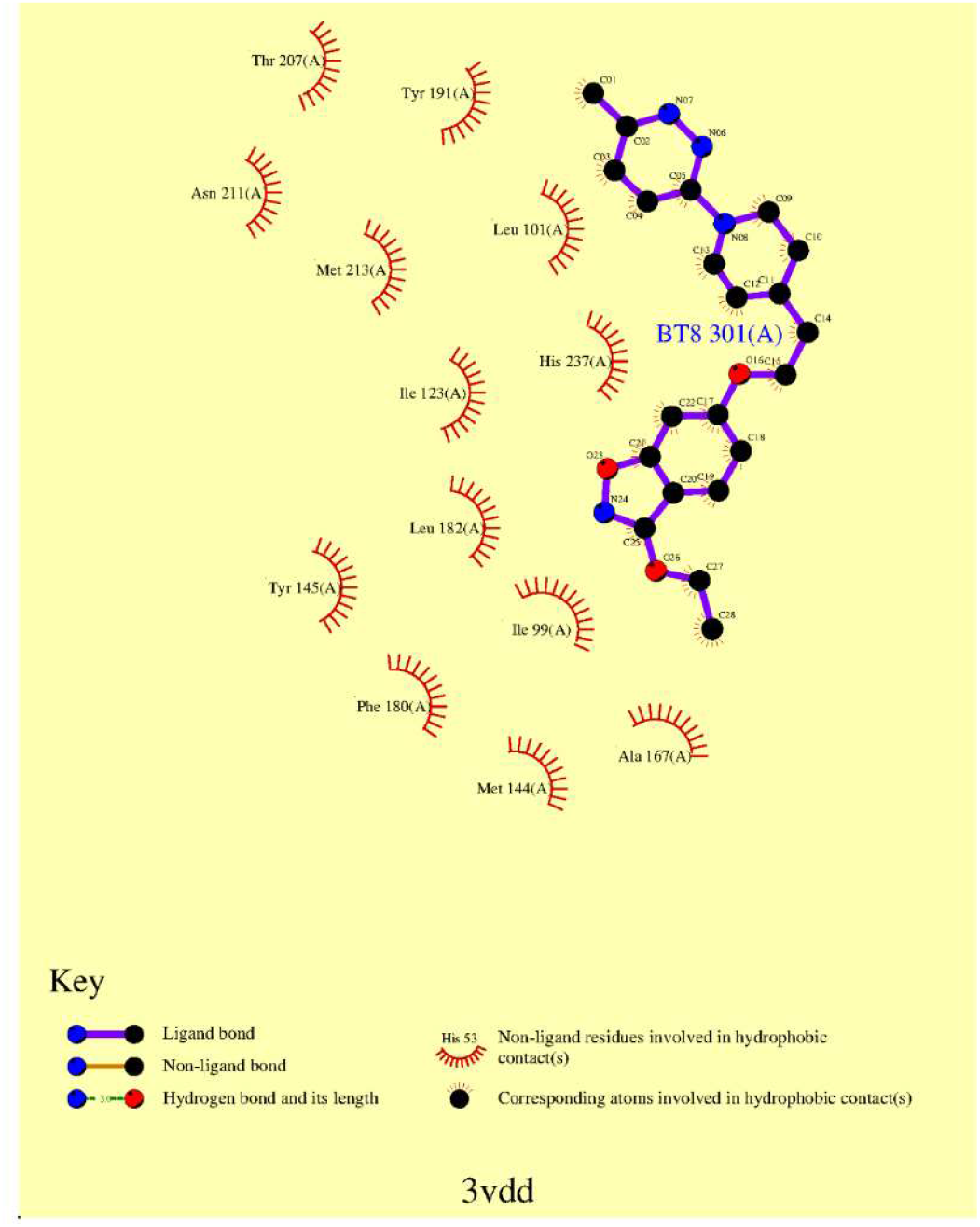
Ligplot map of interactions of the HRV2 capsid with vapendavir in pdb entry 3vdd. M213 makes van der Waals contact with the inhibitor.

**Figure 5.**
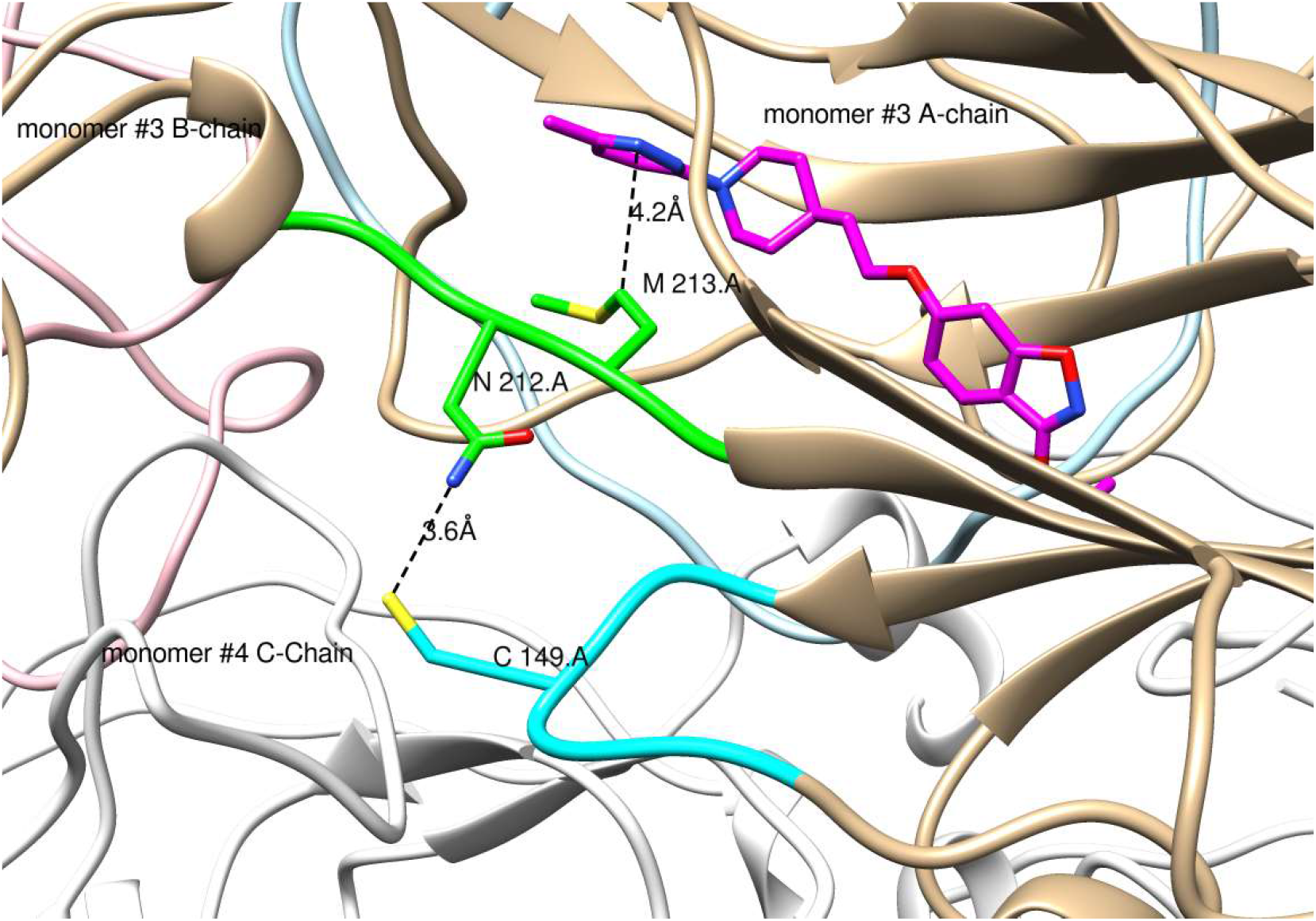
Representative structure of the G149 mutant extracted from the MD trajectory showing the interaction network from the C149 mutation to the inhibitor (pink carbons) binding site in monomer #3 of the pentamer. Intermediate residues involved are N212 and M213. Ribbon colors: Monomer #3 chain A: tan; monomer #3 chain B: pink; monomer #3 chain C: light blue; monomer #3 210-214.a: green; monomer #3 147-152.a cyan; all other ribbons: white. The mutation G149C in monomer #3 A-chain is at the interface of monomer #3 A-chain and monomer #4 C-chain. Image created by UCSF Chimera[31].

## Discussion

We here studied the resistance profile to vapendavir of different enteroviruses. In line with earlier reports, resistant mutants were readily selected in cell culture for all 4 viruses used in the study. The isolates all carried the mutations in the VP1 protein, where the known binding pocket for capsid binders is located. Furthermore, all the isolates proved resistant to other capsid binders confirming the same mechanism of action. Interestingly, we identified two types of mutations conferring the resistant phenotype: i.e. in amino acids lining the binding pocket but also amino acids outside the pocket. Mutants outside the pocket have also been reported in hRV14 isolates resistant to WIN 52084 [32]. However, the mutations identified in hRV2 and EV-D68 are localized in a different loop of the VP1 protein outside the pocket than reported in that study.

We identified pocket mutations in PV1 (Sabin), EV-D68 and hRV14. The pocket mutations impair the binding of WIN-like compounds and thus abolish their antiviral activity. The I194F mutation in poliovirus, as identified here, has been shown to confer resistance to other capsid binders, i.e. H1PVAT and V-073 [28,35]. According to modelling studies, the I194 amino acid is important for the interaction of the capsid binder with the pocket and the mutation in this position can have an effect on compound binding [28]. The pocket mutation at position C199 (located at the pocket entrance) in hRV14 was identified previously in a pleconaril-resistant isolate [21] and in WIN 52084-resistant isolates [32]. A mutation at the entrance to the pocket has been reported as well in a CVB3 isolate resistant to pirodavir [36]. In that case, molecular modelling suggested that mutations in residue I1207 to K or R are preventing the compound from entering the canyon.

Previously reported pleconaril-resistant hRV2 isolates carry the I99F mutation located in the binding pocket [21]. The EV68 mutation V69A in a pleconaril-resistant variant is also located in the pocket [20]. The pocket mutation A156T identified here in vapendavir-resistant isolate of EV-D68 corresponds to the A150T/V mutation reported in hRV14 pleconaril-resistant [21]. However, we here observed mutations in hRV2 and EV-D68 that are located outside the drug binding pocket. Interestingly, the hRV2 resistant isolate had a particular vapendavir-dependent phenotype; its infectivity is dependent on the presence of vapendavir and its replication is impaired in the absence of the drug. On the other hand, the resistant EV-D68 variants carrying the K167E mutation in the same loop of VP1 protein could replicate independently of vapendavir, and their phenotype was similar to that of the isolates with the mutations inside the binding pocket. It is likely that the mutations in the pocket A156T and M252L are primary responsible for the resistant phenotype of EV-D68, since the glutamic acid at the same position as K167E in CU70 strain (K155 in the prototype Fermon strain) is naturally present in several EV-D68 strains, that are inhibited by capsid binders (e.g. 4311000742, 4310900947 and 4310902042). Moreover, this amino acid is present in the 4310901348 strain, which is susceptible to pleconaril and pirodavir, but not vapendavir inhibition [20]. This suggests that this particular residue is not crucial for the antiviral activity of capsid binders. The K167E amino acid change was reported as a secondary mutation in one of the pleconaril-resistant EV-D68 isolates [20]. Taken together, the K167E substitution observed here in the EV-D68 resistant isolates could be beneficial for virus fitness, but is unlikely to underlie the resistant phenotype.

The dependency of picornavirus replication on antivirals has been reported before. For example a Gua-HCl dependent PV1 virus with mutations in 2C protein can only replicate in the presence of Gua-HCl [37]. The dependence of echovirus on rhodanin is attributed to the F53Y mutation in the structural protein VP4 [38]. Dependency of PV3 (Sabin) on WIN 51711 was attributed to mutations in the inside lining of the capsid [39,40].

The hRV2_C3 mutant virus identified here is dependent on the inhibitor for efficient dissemination. Since the isolate has the same replication kinetics as the WT virus and is able to release viral progeny (as shown by detection of the viral RNA in the supernatant of transfected cultures), we suggest that vapendavir binding may be required for the proper conformation of the mutant virus particle allowing it to efficiently bind/enter the cells.

The lack of thermostabilizing effect of vapendavir in the mutant hRV2 suggests that the binding of the compound to the mutant virus differs from the binding to the WT virus. On the other hand, the mutant virus still retains its infectivity at 52°C, whereas the WT variant does not. Since hRV2_C3 stocks were prepared in the presence of the compound it cannot be excluded that the stability of the mutant isolate at 52°C is due to some remaining vapendavir (carry-over effect). Extreme thermolability of WIN 51711-dependent poliovirus mutant particles has been reported [39,40]. WIN_51711 was shown to stabilize the infectious particles without affecting their infectivity. However, the level of resistance level and replication efficiency reported in that study is apparently less pronounced than with our hRV2_C3 isolate.

Our molecular simulation data do not reveal an impairment of vapendavir binding to the mutated VP1 protein of hRV2. This can support the hypothesis that the compound is still capable of binding and stabilizing the mutant hRV2 virus. However, it is possible that the effect of the mutation in simulations may only appear after much longer simulation times. Passaging of hRV2_C3 or EV-A71 BrCr (G159C) without antiviral pressure results in a rapid reverting to wild-type, indicating that the mutation has an infavourable effect on the virus.

In conclusion, vapendavir results, depending on the virus that it inhibits, in various drug-resistant geno- and phenotypes. Like other capsid binders it results rapidly in the selection of drug-resistant variants [12,41,42]. The combination of capsid binders with other entero-/rhinovirus inhibitors that have a different mechanism of antiviral activity (GPEI) may be worth considering.

## Acknowledgements

We would like to thank Vaxart Inc, Dr. Vernachio and Dr. Tucker for providing vapendavir for this study. This project has received funding from the European Union’s Horizon 2020 research and innovation programme under the Marie Sklodowska-Curie grant agreement No 642434. L.S. was funded by the China Scholarship Council (CSC) grant 201403250056.

