## Supplementary figures and tables for "Comparative analysis of the molecular mechanism of resistance to vapendavir across a panel of picornavirus species"

### Supplementary information

**Table S1. Reversed-engineered PV1 and hRV14 viruses confirm the drug-resistant phenotype.**

|  |  | EC_50_ µM |
| --- | --- | --- |
| Virus | **Genotype VP1** | **Vapendavir** |
| hRV14 | WT | 0.09 ± 0.01 |
|  | C199R | >10 |
|  | C199Y | >10 |
| PV1_Sabin | WT | 2.6 ± 0.04 |
|  | I194F | >10 |

**Table S2: Molecular dynamics simulation summary**

| **system** | **Wt + vapendavir** | **G149C + vapendavir** | **Wt unliganded** | **G149C unliganded** |
| --- | --- | --- | --- | --- |
| configuration | Pentamer | Pentamer | pentamer | Pentamer |
| Total atoms | 370819 | 372366 | 372076 | 372099 |
| Counterions (Na+) | 40 | 40 | 40 | 40 |
| Water molecules (periodic truncated octahedral box) | 102628 | 103137 | 103137 | 103138 |
| Temperature | 300 K | 300 K | 300 K | 300 K |
| Pressure | 1 atm | 1 atm | 1 atm | 1 atm |
| Time step | 2 fs | 2 fs | 2 fs | 2 fs |
| Simulated time | 60 ns | 60 ns | 60 ns | 60 ns |
| Ensemble | NPT | NPT | NPT | NPT |
| Electrostatic interactions | PME | PME | PME | PME |
| X-H bonds restraints | Shake | Shake | shake | Shake |
| Protein parameters | ff14SB | ff14SB | ff14SB | ff14SB |
| Vapendavir parameters | Gaff2 | Gaff2 | Gaff2 | Gaff2 |

**Table S3. Distances from cys149.a@SG to asn212.a@OD1 and asn212.a@ND2 calculated from the 60 ns trajectory of the unliganded mutant simulation.**

| **Asn212.a@** | **Average distance (Å)** | **Standard deviation (**$\boldsymbol{Å}$**)** | **Ymin (**$\boldsymbol{Å}$**)** | **Ymax (**$\boldsymbol{Å}$**)** | **N (<4Å) of 15000** | **Monomer #** |
| --- | --- | --- | --- | --- | --- | --- |
| OD1 | 5.975 | 1.353 | 2.931 | 12.53 | 1531 | #1 |
| ND2 | 5.979 | 1.916 | 2.964 | 12.59 | 2110 | #1 |
| OD1 | 5.195 | 1.11 | 2.9 | 11.68 | 1771 | #2 |
| ND2 | 4.067 | 0.9937 | 2.886 | 11.27 | 9671 | #2 |
| OD1 | 5.87 | 1.222 | 2.891 | 10.85 | 831 | #3 |
| ND2 | 6.079 | 1.813 | 2.76 | 11.47 | 2479 | #3 |
| OD1 | 5.783 | 1.63 | 2.901 | 11.99 | 1902 | #4 |
| ND2 | 5.039 | 1.69 | 2.911 | 11.49 | 5295 | #4 |
| OD1 | 4.857 | 1.032 | 2.936 | 11.43 | 3206 | #5 |
| ND2 | 4.693 | 1.421 | 2.862 | 11.12 | 6641 | #5 |

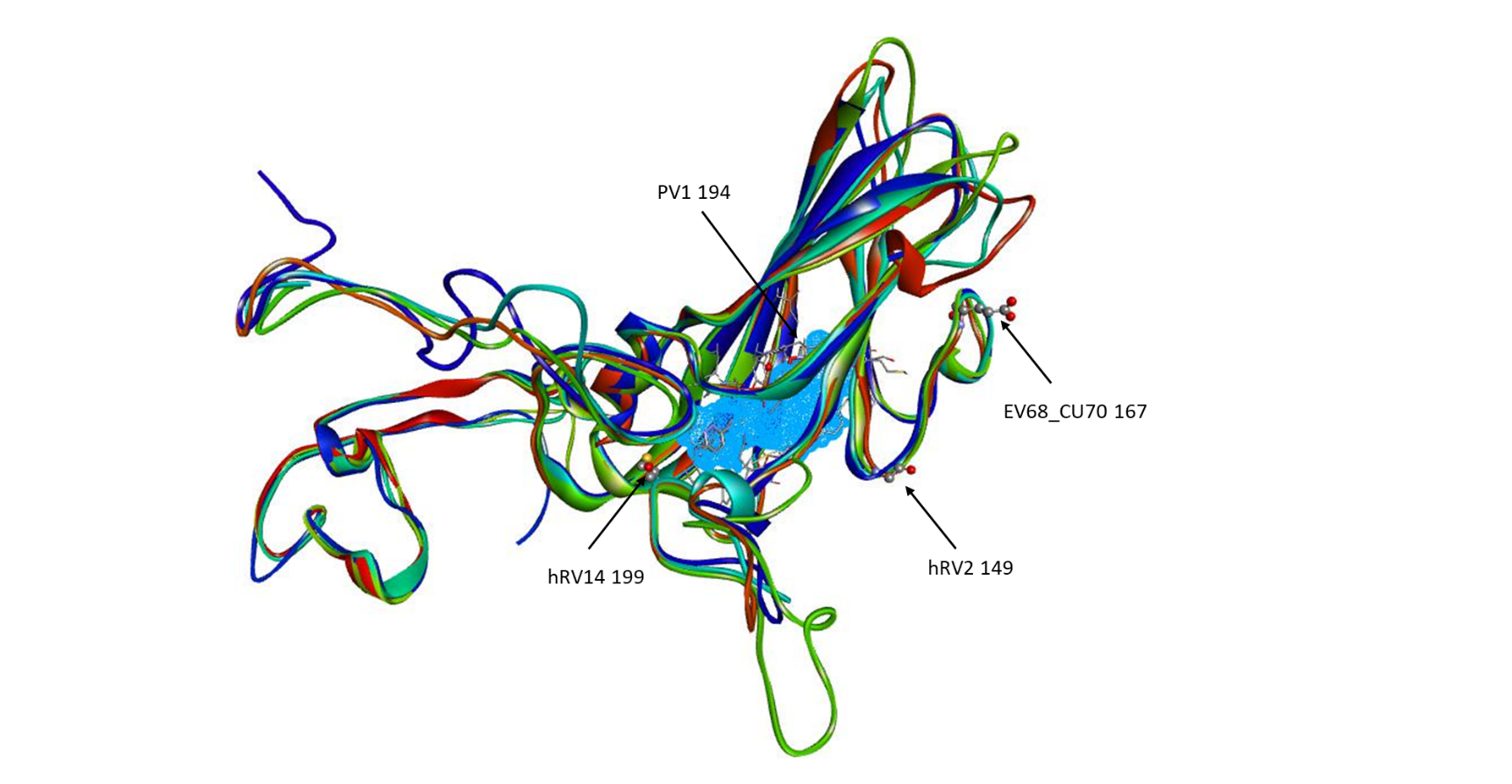

**Figure S1. VP1 structures of viruses used in this study aligned to hRV2 VP1 in complex with vapendavir.** hRV14 (4hrv) in red, PV1 (1vbd) in green, hRV2 (3vdd) in turquoise, EV-D68 (4wm8) in blue. Mutated residues in sticks and balls. The structure alignment performed with UCSF Chimera.

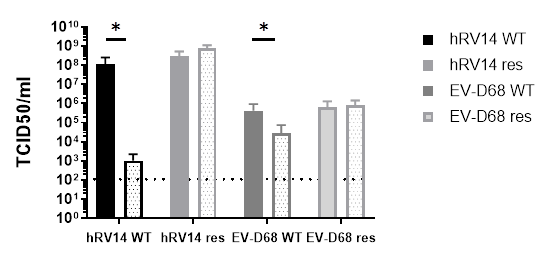

**Figure S2. hRV14 and EV-D68 mutants do not show vapendavir-dependent phenotype.** Infectious virus load measured by end-point titration in presence (solid filled bars) or absence (dotted bars) of vapendavir. The graph represents mean of 3 independent experiments. * p<0.05, t-test.

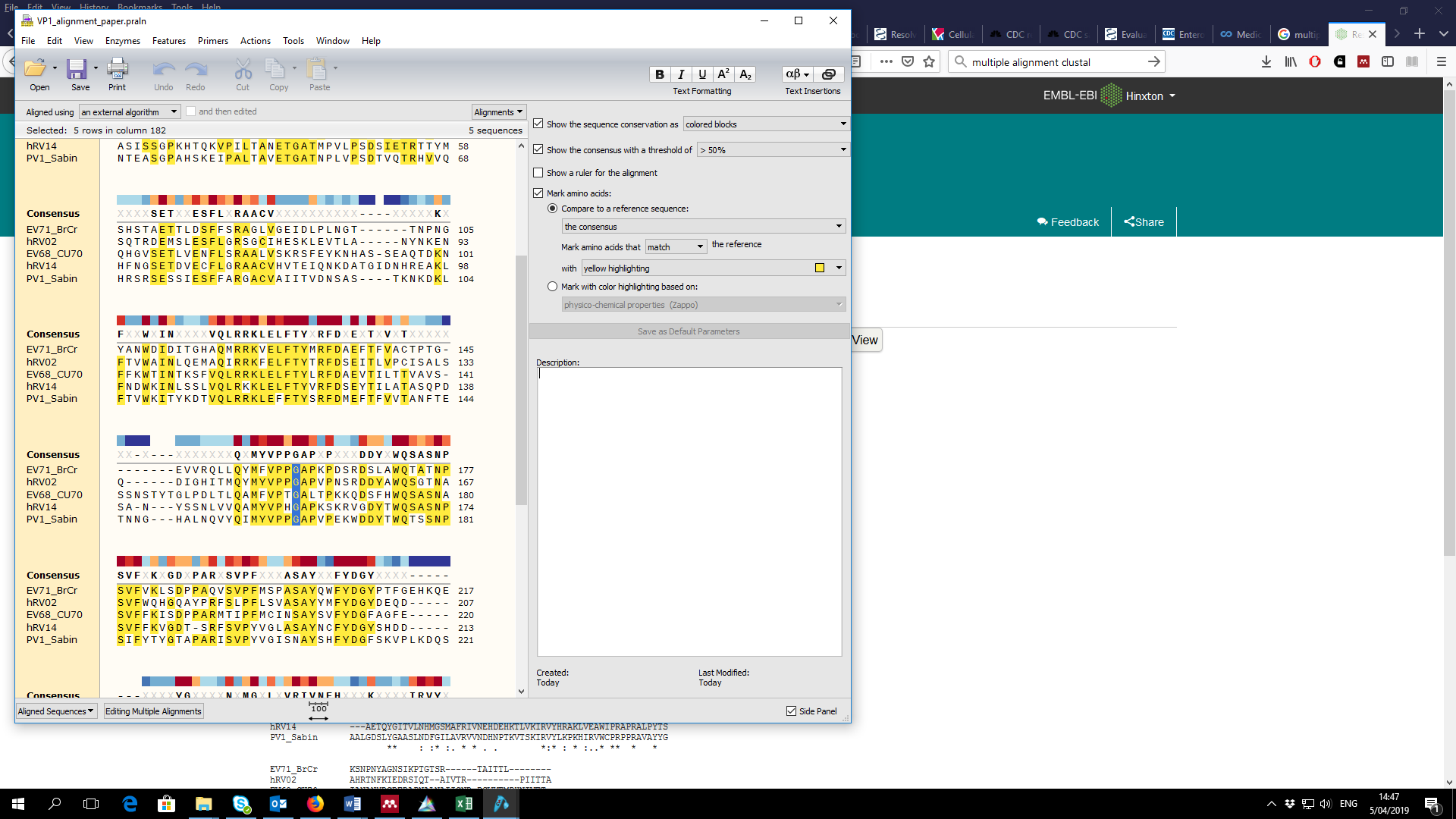

**Figure S3. Alignment of VP1 protein AA sequence in different enterovirus species**
